# Experimental Evolution of the Thermal Performance Curve

**DOI:** 10.1101/2025.09.23.677994

**Authors:** Giacomo Zilio, Iain R. Moodie, Sarthak P. Malusare, Marie-Ange Devillez, Justina Givens, Claire Gougat-Barbera, Emanuel A. Fronhofer

**Author notes:** **Correspondence** Details Emanuel A. Fronhofer, Institut des Sciences de l’Evolution de Montpellier, UMR5554, Université de Montpellier, CC065, Place E. Bataillon, 34095 Montpellier Cedex 5, France. shared first authorship (Giacomo Zilio, Iain R. Moodie, Sarthak P. Malusare).

## Abstract

The thermal performance curve (TPC) of an organism captures how population growth depends on temperature. When populations experience increased temperatures, such as during global climate change, one prediction is that their TPC can evolve to accommodate the new environmental temperature. Although studies on TPC evolution have mostly focused on modifications in population growth rates, TPC evolution can be strongly trait-dependent and require a multi-trait analysis. Thus, if and how the entire TPC across a multitude of traits evolves to change its shape in response to increased temperatures remains debated. Here, we empirically tested how the TPCs of multiple demographic, life-history and movement traits can evolve by selecting four freshwater protist species at increased temperatures starting from clonal populations. After one year of selection, populations showed a signature of evolutionary responses to the highest selection temperatures in different traits depending on the species. Particularly, we found consistent evolutionary reductions in body size in the three species having the largest cells and evolved changes in movement behaviour in all species. In contrast, we overall observed little modifications in population growth rate and in the corresponding TPC shape. These results suggest that adaptation, via evolution of TPCs, might involve the concurrent evolution of several traits. However, this may be species-specific and difficult from de-novo mutation alone, suggesting that natural populations that do not have sufficient standing genetic variation might have to be reliant on other means of mitigating the effects of climate change, such as dispersal.

## Introduction

In ecology and evolutionary biology we strive to understand how populations respond to novel environments (Merilä & Hendry, 2014; Mouquet *et al*., 2015; Urban *et al*., 2016). Populations that experience substantial environmental stress, such as global climate change, must either adapt to the stress or disperse away from the stress, otherwise they will face population decline and an increased risk of extinction (Thomas *et al*., 2004; Parmesan, 2006; Gienapp *et al*., 2008; Urban *et al*., 2024).

One central aspect of global climate change is increasing temperatures and global warming (Hansen *et al*., 2006). Temperature is an environmental variable that affects nearly all biological processes, from the molecular and biochemical dynamics of enzymes (Gillooly *et al*., 2001) to population growth rates, spatial patterns and diversity of populations and communities (Angilletta, 2004; Wiens *et al*., 2006; Kingsolver, 2009).

To be able to predict the effects of global climate change on populations, it is imperative to understand if and how adaptation to increased temperatures can occur, how quickly evolution can act, and if these adaptations are associated with trade-offs or correlated responses more generally that would impact other traits which may limit their effectiveness (Bennett & Lenski, 2007). For instance, advancements in the field of metabolic ecology allow us to predict how ecological rates (e.g., population growth rates, developmental rates) scale with temperature and related factors (e.g., body size) through the use of semi-mechanistic models (Brown *et al*., 2004).

Thermal performance curves (TPCs), which describe the thermal plasticity (or temperature niche) of a trait across a range of temperatures, are a powerful tool to investigate the impact of warming under controlled conditions (Schulte *et al*., 2011; Deutsch *et al*., 2008). Here, we specifically define the temperature niche, or thermal performance curve (TPC), of an organism as the range of body temperatures at which a non-density limited population will show positive population growth (Gvoždík, 2018). The TPC, and the reaction norms of other traits (e.g., developmental rates, fecundity, body size), is typically unimodal in ectotherms, and is often described by the temperature value at which the trait is maximised by (the thermal optimum, *T*_*opt*_), and the range of temperatures at which an organism can respond (the width) (Kingsolver *et al*., 2011; Amarasekare & Savage, 2012; Grimaud *et al*., 2017). Further, the same shape (approx. left skewed unimodal distribution) of the temperature niche appears across a wide-range of ectotherm taxa (Deutsch *et al*., 2008; Kingsolver, 2009; Dell *et al*., 2011), likely due to the highly conserved nature of the physiological and biochemical processes that underlie these reaction norms (Gillooly *et al*., 2001; Have, 2002; Brown *et al*., 2004).

Multiple empirical studies have found that aspects of the TPC can evolve when exposed to selection. Using outdoor mesocosms, Wang *et al*. (2023) found in the crustacean *Daphnia magna* rapid evolution of TPCs for multiple key traits (e.g., survival, fecundity, growth rate), but not traits related to energy gain. Schaum *et al*. (2017) carried out a decade long warming experiment with the phytoplankton *Chlamydomonas reinhardtii* and found that isolates from warmed mesocosms had higher optimal temperatures (*T*_*opt*_) than control isolates, and consequently had higher competitive ability at warmed temperatures than control isolates, a strong indication of adaptation to increased temperature. In a step-wise temperature selection experiment designed to generate thermally adapted lines of the coral photosymbionts *Symbiodinium*, Chakravarti & van Oppen (2018) observed stable adaptive change at the end of their exposure period (41–69 generations), resulting in populations that showed faster growth rates under acute heat stress, and higher photosynthetic efficiencies when compared with control populations, although these results were not found across all replicate lines. Bennett & Lenski (1993) evolved replicate lines of *Escherichia coli* for 2000 generations at three fixed selection temperature (32, 37, 42°C), or at a temperature regime that routinely fluctuated between the two extremes. They then measured the range of temperatures (12–44°C) that a population could maintain itself while undergoing serial dilutions. They found that the width of the temperature niche and extinction risk at high temperature was unaffected by the selection regime, but each selection line showed a pattern of local adaptation, an increase fitness at their selected temperature. In a similar experiment in *E. coli*, but specifically looking for trade-offs to high temperature (40°C) after adaptation to 20°C, Bennett & Lenski (2007) did find evidence of a general trade-off (a decline in mean fitness across all replicates), although they were not universal across replicates. A meta-analysis of the field of thermal selection experiments (Malusare *et al*., 2023) could recently show that TPCs have adaptive potential across ectothermic species. This synthesis clarifies that trade-offs between adaptation to higher temperatures and fitness at lower temperatures can be generally observed across the tree of life. At the same time this study highlights the relative paucity of well-resolved TPCs and the limit of using only two test temperatures, as done by many studies, which hinders interpretation related to shape changes and responses at thermal limits.

Despite these recent advances and broad interest, more work remains to be done to increase our understanding of how thermal stress impacts the evolution of TPCs. Furthermore, many temperature selection studies focus on a single species, which makes comparisons of how species respond to identical selection regimes difficult. A better understanding of the differences between species’ ability to adapt their plastic response to increased temperature could help in predicting the evolution of community composition under climate change, as well as the relative rates of adaptation and extinctions in different environments.

In order to address these challenges, we performed such a multi-species temperature selection experiment. After 1 year of thermal selection at six different temperatures, we empirically tested how TPCs of multiple traits respond to selection across four life-history diverse protist species. All species showed responses to thermal selection and differences in their TPCs compared to the evolutionary control treatment. Detailed responses differed across species, and mainly involved changes in body size and movement rather than population growth rate.

## Materials and Methods

### Study organisms

In this experiment we used four freshwater protist species, spanning a broad range of body sizes and trophic diversity: two genotypes of *Tetrahymena thermophila* (*Tet* ; MT I: SB3539 and MT VII: CU428.2), a small (approx. 50 µm) bacterivorous ciliate commonly used in evolutionary and ecological microcosm experiments (Collins, 2012; Altermatt *et al*., 2015; de Melo *et al*., 2020; Moerman *et al*., 2020a,b), *Paramecium caudatum* (*Para*), a large (approx. 330 µm) widely studied bacterivorous ciliate (Magalon *et al*., 2010; Nørgaard *et al*., 2021; Zilio *et al*., 2023a), *Euglena gracilis* (*Eug*), a small (20-100 µm) mixotroph capable of both photosynthesis and phagocytosis (Altermatt *et al*., 2015; Harvey *et al*., 2017), and *Blepharisma* sp. (*Ble*), a large (approx. 200 µm) omnivore that feeds on bacteria and smaller ciliates (Giese, 1938; Saade *et al*., 2022; Tan *et al*., 2021).

These species were chosen due to their wide diversity in life-histories and their potential for studying biotic interactions among species in mixed species microcosms (Carrara *et al*., 2015). Populations were cultured using medium made from dehydrated organically grown salad and Volvic mineral water (1 g of salad in 1.6 L of water), autoclaved and then inoculated with the bacterium *Serratia fonticola* at 10% maximum density (tenfold dilution of one week old culture at 20°C is a food source for the protist species.

### Selection protocol

Temperature selection lines (20, 25, 30, 33, 36 and 39°C) were generated for each species/strain using a ratchet protocol (modified from Huertas *et al*. 2011). Each selection temperature included three replicates that evolved independently, for a total of 90 selection lines (5 species *×* 6 temperatures *×* 3 replicates). This method strikes a balance between strong selection, through the step-wise increase in temperature, and maintaining high population sizes, to maximise the number of spontaneous mutations conferring adaptation to the increase in temperature.

In brief, three biological replicate populations of each organism were established from clonal lines derived from laboratory stock cultures at 20°C. After six weeks, 10 mL samples were taken and established in 90 mL of 10% bacterized salad medium at the next higher selection temperature. If a population showed positive population growth after six weeks at a selection temperature, a sample was ratcheted to the next selection temperature (Fig. S1). If not, it was evaluated again after another six weeks. If an extinction occurred, a new population was established from the previous selection temperature. Selection lines were stored in temperature controlled incubators (randomised twice per week to avoid incubator effects) with a 12:12 light:dark photoperiod. To maintain excess food and prevent population growth rates declining, 10 mL from each selection line was regularly transferred into 90 mL of 10% bacterized salad medium (Tet I and Tet VII: 7 days, Para and Ble: 14 days, Eug: 21 days; the different timing reflects overall population growth rate differences).

As a reference, we used the 20°C selection line. We did this, as opposed to using ancestral measurements, to minimise time artefacts in the data and to ensure that the only difference between the reference selection line and other selection lines was the selection temperature. The selection regime may have introduced other unintended selection pressures (e.g., during weekly transfers), and ensuring that these were identical across all lines, including the reference, allowed for direct comparisons regarding the effects of temperature within and between selection lines.

### Assay Protocol

The goal of the assay was to measure population growth, as well as other traits (see below), of each selection line at ten different temperatures (10, 15, 20, 25, 30, 33, 36, 39 and 40°C) over a period of three weeks. The three replicates were measured in three temporal blocks. Prior to beginning the temperature assay, we kept the selection lines in a 20°C common garden phase to minimise parental effects, adjusting approximately the time proportional to each species’ generation times (*Tet* I and *Tet* VII: three days; *Para* and *Ble*: five days; *Eug:* seven days). To establish the assay lines, we added 200 µL (20 µL for *Tet* I and *Tet* VII) of selection line protist culture to sterile plastic vials containing 20 mL of 10% bacterized salad medium and stored them in temperature controlled incubators (randomised twice per week) with a 12:12 light:dark photoperiod. Three times per week, we adjusted the volumes of each assay line vial to 18 mL and added 2 mL of 10% bacterialised salad medium to ensure sufficient food availability.

At each time-point, we placed 350 *µ*L samples from assay line vials into 24-well clear bottom microplates (Porvair Sciences, UK). We imaged each sample using a microscope plate reader (Cytation 5, BioTek) for 10 seconds (150 frames). The video files were then analysed using the ‘BeMoVi’ R package (Pennekamp *et al*., 2015) to extract density, cell size data and swimming movement data (for exact settings and modifications see https://github.com/efronhofer/analysis_script_bemovi/tree/ffmpeg_overlays). The whole procedure was firstly used to determine the ancestral TPC and traits of each species, and it was then repeated after one and two years of selection to determine evolutionary changes. During the 2 years, 18 lines across different species and selection temperature went extinct, leaving us with 83 lines for year 1, and 72 for year 2 (Table S1). Data from year 1 and 2 show no overall different pattern (Fig. S2), so we therefore focus on data from year 1.

### Statistical analyses

All statistical analyses were performed using R v4.2.0 (R Core Team, 2022) and the packages ‘*rstan*’ v2.21.5 (Stan Development Team, 2022) and ‘*brms*’ v2.17.0 (Bürkner, 2017).

### Data check and cleaning

In order to facilitate the statistical analysis, we first proceeded to visually inspect each population time series collected during the assays and clean data to exclude potential outliers and cases in which populations did not grow. Upon inspection of the video files and ‘BeMoVi’ overlays, we found that some outliers in population densities were caused by the plate reader having failed to auto-focus correctly. In order to remove such outliers systematically, we first fitted LOESS splines using the loess() function in R to the data and removed any point that deviated too far (3 times greater, respectively lesser) from the spline fit. We then excluded any populations with less than 7 data points in their time series, reducing the initial 830 populations to 766.

### Population growth rates and other traits

To extract estimates of population growth rates (*r*_0_), we fitted Bayesian linear models to the population density estimates (log-transformed) during the exponential phase. The exponential phase was identified using visual inspection of all growth curves. We used an MCMC chain length of 30,000 iterations (15,000 warmup iterations) and vaguely informative priors, with the intercepts and slope parameters following a normal distribution and half-normal distribution respectively, with mean 0 and standard deviation 1. The adapt delta parameter was set to 0.999 to eliminate post-warmup divergent transitions. The estimated slope parameters corresponded to the population growth rates. Measures of other traits included the major and minor cell axes (proxies for body size), bioarea (proxy for biomass), as well as swimming speed and tortuosity (s.d. of the turning angle distribution). These were averaged over the exponential growth phase selected. We additionally considered the time point at which populations attained their maximum density as a proxy for equilibrium density.

### TPC fitting

To explore how the TPC varies after selection at increased temperatures, we fitted thermal performance curves to the *r*_0_ estimates obtained as described above. We use a high temperature inactivation version of the Sharpe-Schoolfield equation (Schoolfield *et al*., 1981), a commonly used semi-mechanistic model that extends the Arrhenius equation to incorporate a decline in population growth rates beyond an optimum value. The model is given by the equation

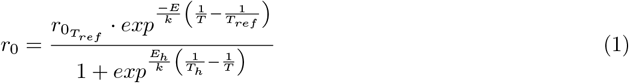

where *T* is temperature in Kelvin (*K*), *k* is the Boltzmann constant (8.617 *×* 10^−5^*eV K*^−1^), *E* is the average activation energy (eV), *E*_*h*_ is the high temperature deactivation energy, *T*_*h*_ is the temperature at which growth rates are 50% decreased due to high temperature and *r*_0*ref*_ is the *r*_0_ value at a reference temperature (here, 20°C). The optimum value (*T*_*opt*_) of Eq. 1 is given by

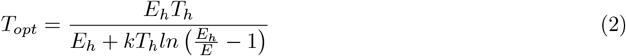

and the maximal growth rate (*r*_*max*_) can be calculated by substituting *T*_*opt*_ into Eq. 1.

We again made use of a Bayesian statistical model to fit the TPC function to the *r*_0_ estimates, which allowed for the propagation of the error associated with *r*_0_ estimates throughout the analysis. We estimated parameter values for 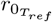, *E, E*_*h*_ and *T*_*h*_ using the ‘*brms*’ and ‘*rstan*’ R packages (for R code, see Supplementary Methods). As before, we used priors that were informative, making use of published values to inform mean and variance estimates of *E* and *E*_*h*_, and using realistic, but broad priors for *T*_*h*_. The assumption of a universal temperature dependence (i.e., that values of *E* are highly constrained by thermodynamics), is often called into question (Clarke, 2004; Clarke & Fraser, 2004; Clarke, 2006; Gillooly *et al*., 2006), hence we used wider priors for *E* than those directly predicted from metabolic theory of ecology. The only prior that differed between species was 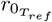 (for full information on the priors used, see Table S2). We included a measure of the error on each *r*_0_ estimate by including each estimate as a normal distribution with a mean and standard deviation taken from the posterior distribution of the previous analysis (normal distribution on the log scale). For each selection temperature of each species of each replicate, we ran chains with 10000 iterations (warmup=5000 iterations) (warmup=0, adapt delta=0.99).

### TPC parameters

The TPC parameter posteriors 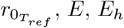 and *T*_*h*_ directly estimated from the TPC fitting were scaled and centred around their means. These were analysed using multi-variate multilevel models (warmup = 10,000 iterations, chain = 20,000 iterations) with a normal error structure. We conducted two main analyses for each species. Firstly, to assess potential TPC responses to laboratory conditions we compared the intercept model to the one including the ancestral populations and the evolutionary control lines at 20°C as explanatory factors (i.e., evolutionary conditions). For model comparison and weight we used the Watanabe-Akaike Information Criterion (WAIC; Watanabe & Opper, 2010), a generalized version of the Akaike Information Criterion (Gelman *et al*., 2014). Secondly, we applied the same statistical procedure to test for TPC differences across the selection temperatures, from the evolutionary control at 20°C to the highest-temperature evolutionary treatment at 39°C. In this second analysis we considered selection temperature as a continuous response variable, and we further included replicates as a random factor. In all models we adopted the default vaguely informative priors of the ‘brms’ package. Additionally, we used the obtained 4-parameter matrix for complementary graphical inspection by means of principal component analysis (PCA).

For the calculated parameters *T*_*opt*_ and *r*_*max*_ we proceeded similarly. We conducted 2 separate main analyses for each of the 5 species using univariate multilevel models (warmup = 10,000 iterations, chain = 20,000 iterations) with a normal error structure. In one analysis we compared WAIC of the intercept to the evolutionary conditions model, and in the other the intercept to the selection temperature model. We also propagated the error associated with *T*_*opt*_ and *r*_*max*_ estimate calculation. In some cases, when analysing *r*_*max*_, the inclusion of the selection line random effect in the selection temperature models produced few divergent transitions (*<* 0.01%) even after applying guidelines for corrections. Following Stan Development Team diagnostics, careful inspections revealed no pattern in their locations, and the models had ideal convergence (Rhat = 1.00) and effective sample size (ESS) values. Further, the results were qualitatively comparable to models without random terms (and divergent transitions). We therefore decided to maintain the models with the random effect structure.

### TPC for phenotypic traits

We applied the same statistical approach as for the TPC parameters described above. For each species, we run Bayesian multi-variate models (warmup = 10,000 iterations, chain = 20,000 iterations) on 7 traits, using normal error structure and model comparison. The phenotypic traits were *r*_0_, swimming speed and tortuosity, major and minor cell axis, bioarea and maximum density, all of which were scaled and centred prior to the 4 main analyses. We compared ancestral and evolutionary control lines (two factors variable) assayed at their own selection regime (20°C) and at the other temperatures. In the other two analyses, we similarly compared evolved lines assayed at their own selection treatment or in the other temperatures. We considered selection temperature as a continuous variable and we also included a selection line random effect. The few cases of divergent transitions were carefully treated as previously described.

## Results

### TPC parameters

For all species, we observed TPC changes in response to laboratory conditions and handling (evolutionary conditions: WAIC weight *>* 0.96; Table S3 and S4). The evolutionary control lines maintained and propagated during a year at standard 20°C showed indeed a different combination of niche parameters compared to the ancestral ones (Fig. S3). Thus, the TPC of the evolutionary control at 20°C is the most reliable baseline information to compare the evolutionary impact of the selection temperatures.

We found some signals of TPC evolution after selection to increased temperatures in both *Tetrahymena thermophila* strains (selection temperature: Tet I WAIC = 192.87; SE = 21.45; WAIC weights = 0.95; Tet VII WAIC = 172.7; SE = 19.89; WAIC weights = 0.98), but not across all species (Fig. 1). For *Tetrahymena thermophila* strain VII, selection at the highest temperatures of 36°C and 39°C decreased 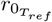 and thus led to a reduction in population growth at the reference 20 C control temperature (slope= -0.096; 95% CI= -0.166, -0.027, Fig. 2). For *Tetrahymena thermophila* strain I, the signal was less clear. Even though no parameter correlation was formally different from 0 (Fig. 2; Table S5), we observed an overall shift in the parameters space for lines selected at the highest temperature of 39°C (Fig. S3 A) similarly to what was found for strain VII (Fig. S3 B). In contrast, the TPCs of *Paramecium caudatum, Blepharisma* sp. and *Euglena gracilis* were unaffected by the selection temperature (Fig. 1; evolutionary conditions: WAIC weight *<* 0.2).

**Figure 1:**
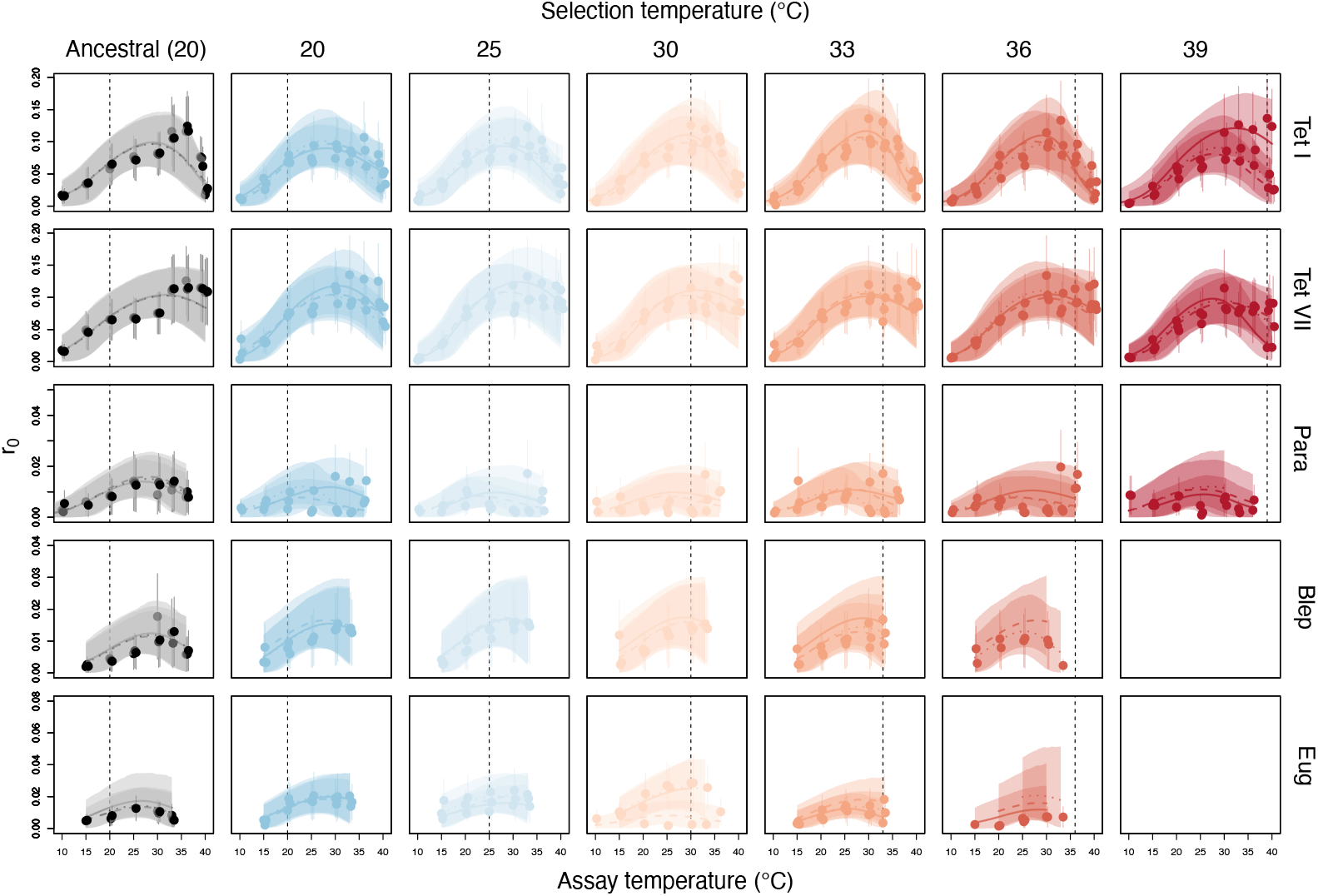
Temperature Performance Curve (TPC) for *r*_0_ (population growth rate) estimates across different assayed temperatures ranging from 10 to 40°C for four protist species subjected to six temperature selection regimes for one year (colour gradient from left to right). The ancestral conditions are represented in grey. Fitted curves and 95% credible intervals, lines and polygons respectively, are derived from a Bayesian implementation of the modified Sharpe-Schoolfield equation (Eq. 1) fitted to *r*_0_ and SE estimates (solid points and bars). The latter were derived from population density estimates recorded over a period of three weeks. Biological replicates are denoted within each panel by a different line type. Dashed vertical lines indicate matching selection and assay temperatures. Tet I and VII are two mating types of *Tetrahymena thermophila*, Para is *Paramecium caudatum*, Blep is *Blepharisma* sp., and Eug is *Euglena gracilis*.

**Figure 2:**
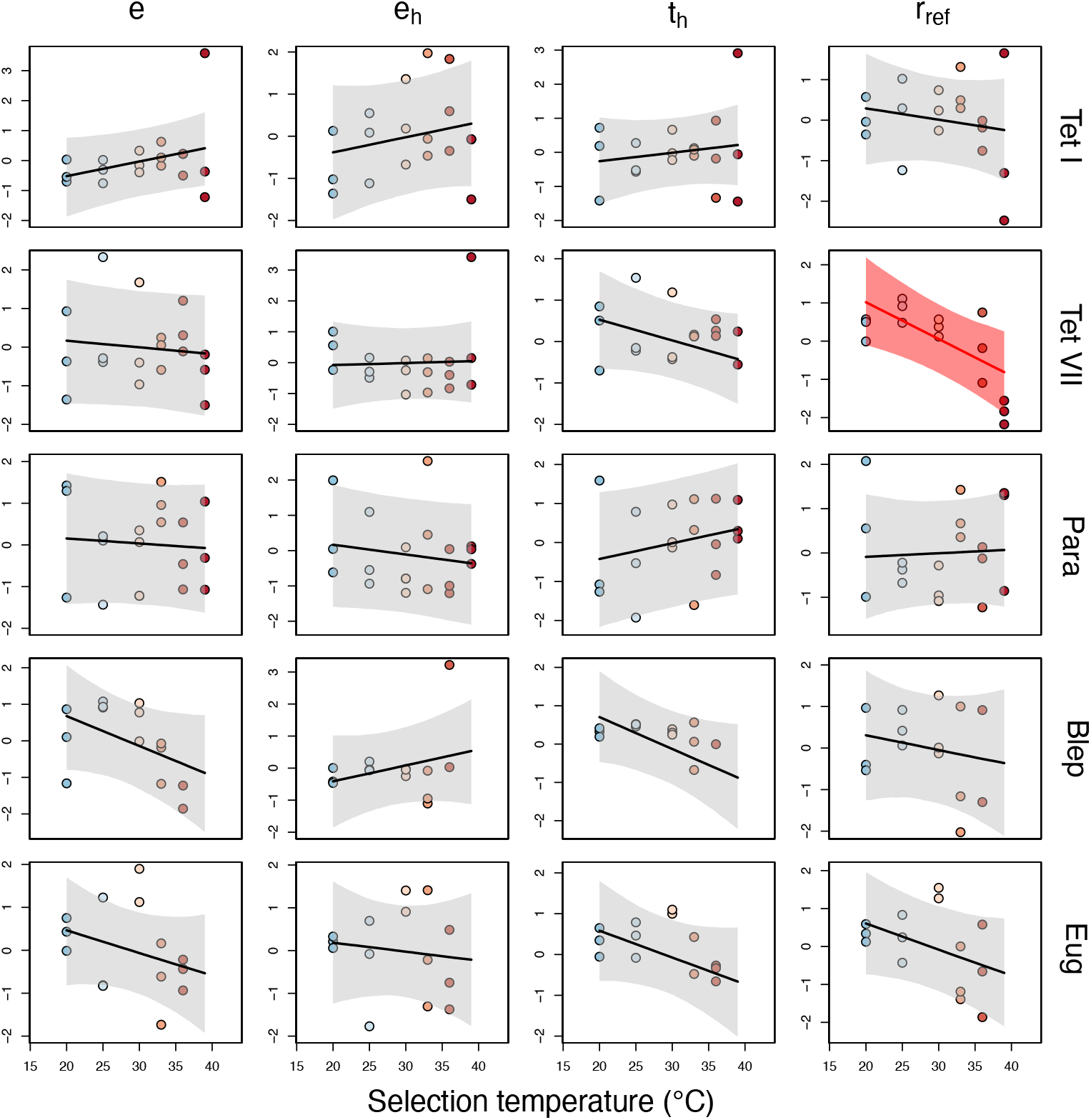
Median model predictions (solid lines) and 95% compatibility interval (shaded areas) obtained from the multivariate analysis of the TPC parameters *e, e*_*h*_, *t*_*h*_, *r*_*ref*_ (niche fitting from Eq. 1). Each point is a standardized niche parameter value corresponding to a biological replicate. The colours are the different selection temperatures experienced during evolution, as also indicated on the x-axis. Model predictions with slope parameter different from 0 are shown in red. Tet I and VII are two mating types of *Tetrahymena thermophila*, Para is *Paramecium caudatum*, Blep is *Blepharisma* sp., and Eug is *Euglena gracilis*.

There were no evolutionary changes in the optimal growth temperature *T*_*opt*_ and maximal growth *r*_*max*_ of all five species compared to the ancestral conditions (*T*_*opt*_: WAIC weight *<* 0.47, Table S6; *r*_*max*_ : WAIC weight *<* 0.43, Table S8). The absence of responses to increased selection temperature was also found across temperatures (*T*_*opt*_: WAIC weight *<* 0.33, Table S7; *r*_*max*_ : WAIC weight *<* 0.47, Table S9). The *r*_*max*_ of *Paramecium caudatum* evolutionary control lines tended to have the opposite trend and be higher than the ancestral conditions, although this remained a weak signal (WAIC = -20.09; SE = 1.18; WAIC weights = 0.57), which was not found when comparing lines across temperatures (WAIC = -78.79; SE = 3.48; WAIC weights = 0.39).

### TPC phenotypic traits

We found changes in phenotypic traits of the evolved control compared to the ancestral lines due to laboratory conditions. This was consistent when measuring them at the same reference 20°C temperature (WAIC weight *>* 0.83) or across assay temperatures, from 10°C to 40°C (WAIC weight = 1; see Table S10 and S11). The evolutionary control lines of all species evolved smaller sizes (reduced minor and major cell axes), but increased their intrinsic growth rate (*r*_0_) and reached higher maximum density, independently of the assayed temperatures (left vs. right column in Fig. S4). *Paramecium caudatum* was somewhat different: In response to laboratory conditions it evolved reduced cell size as the other species, whilst decreasing maximum density compared to its ancestral lines (Fig. S4 E, F). When these evolved lines were tested across temperatures they further exhibited reduced growth rate (Fig. S4 F).

As for the TPC above, we considered the evolutionary control lines as a baseline for additional comparisons. After selection to increased temperatures, populations rarely showed adaptation to their selection temperature (Fig. 3), with notable exceptions (Table S12). We found direct responses to selection (own temperature) only in Tet I (WAIC = 310.78; SE = 14.72; WAIC weight = 1) and *Blepharisma* sp. (WAIC = 239.69; SE = 7.42; WAIC weight = 0.99), particularly at their highest selection temperature, 39°C and 33°C respectively (Fig. 3 A,G).

**Figure 3:**
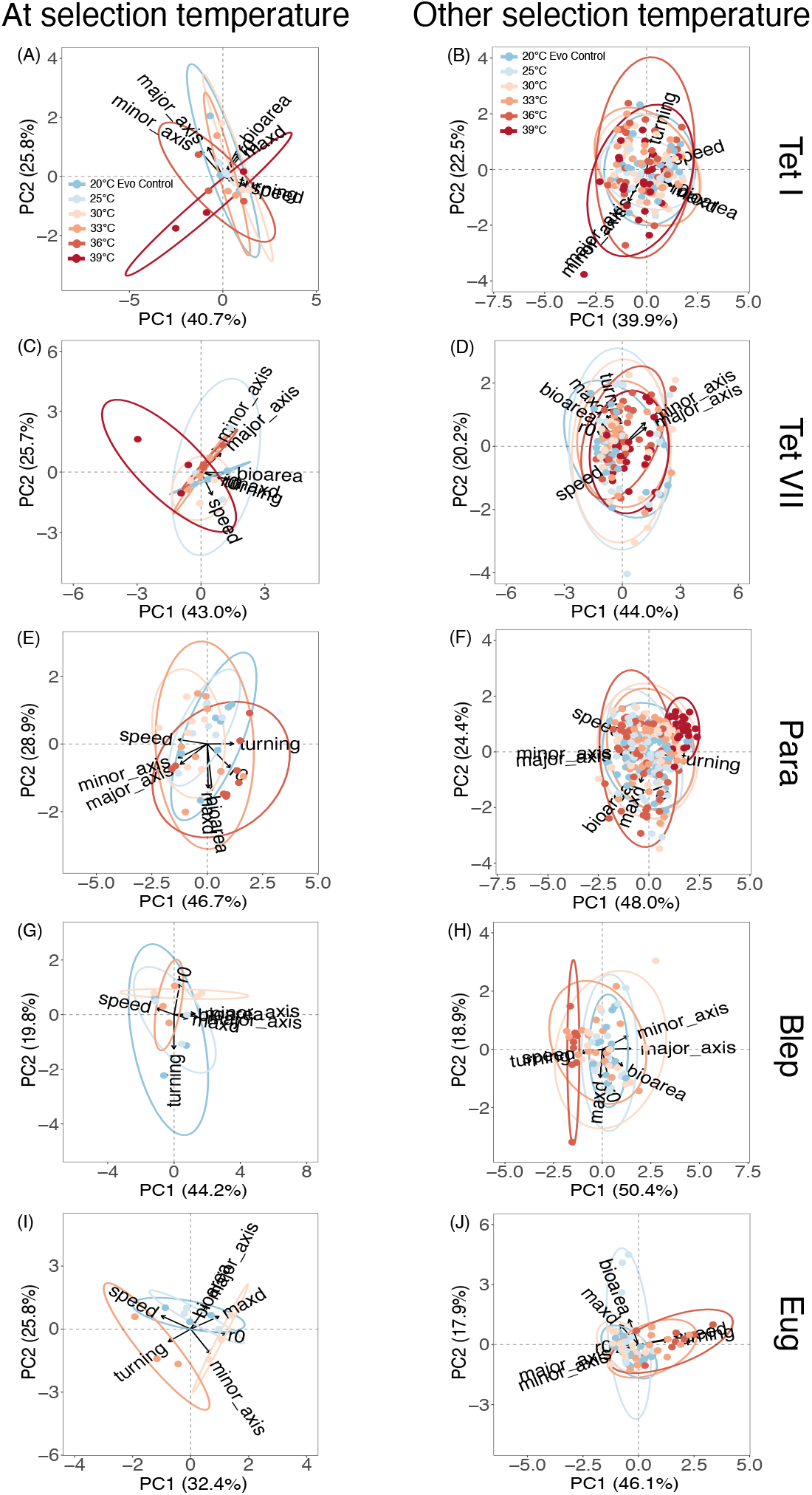
PCA with 95% confidence ellipsis probability of the 7 phenotypic traits measured at their own selection temperatures (A, C, E, G, I) or at others (B, D, F, H, J). Different colours show the selection temperature during the evolutionary experiment. Each of points represents the average phenotypic value in multi-variate space, with arrows indicating the direction of trait variation. Tet I and VII are two mating types of *Tetrahymena thermophila*, Para is *Paramecium caudatum*, Blep is *Blepharisma* sp., and Eug is *Euglena gracilis*.

Note that for *Blepharisma* sp., lines selected at 36°C did not grow at their own selection temperature, and lines at 39°C were not available overall (Table S1). Despite the signal remaining weaker, we also found a trend for direct responses to selection in Tet VII (ΔWAIC = 0.57), which was confirmed by inspecting the model output (Table S13). Tet I and Tet VII reached lower maximum density after selection at higher temperatures (Fig. 3 A,C; Tet I slope= -0.121, 95% CI=-0.169; -0.072; Tet VII slope= -0.074, 95% CI=-0.146; -0.001). In addition, Tet VII decreased its bioarea (slope= -0.072, 95% CI=-0.143; -0.001) and became a slower swimmer (slope= -0.082, 95% CI=-0.157; -0.006). We observed the same trend for trait changes in bioarea and swimming speed in Tet I (Table S13). *Blepharisma* sp. evolved at 33°C and assayed at the same selection temperature increased its intrinsic growth rate (Fig. 3 I, slope= 0.126, 95% CI=-0.001; 0.249). We detected no clear direct responses to selection in *Paramecium caudatum* and *Euglena gracilis* (Intercept: WAIC weight *>* 0.89). However, when considering the whole TPC, i.e., phenotypic traits measured across temperature after selection, all species exhibited evolutionary responses (Table S14; Selection temperature: WAIC weight *>* 0.99). We found stronger and consistent patterns for *Paramecium caudatum, Blepharisma* sp. and *Euglena gracilis* (Fig. 4, Table S15). Namely, selection at the highest temperature led to smaller body size and impacted mobility traits in all 3 species (Fig. 3–4). The swimming behaviour of the Tet VII strain was also affected by selection at higher temperatures.

**Figure 4:**
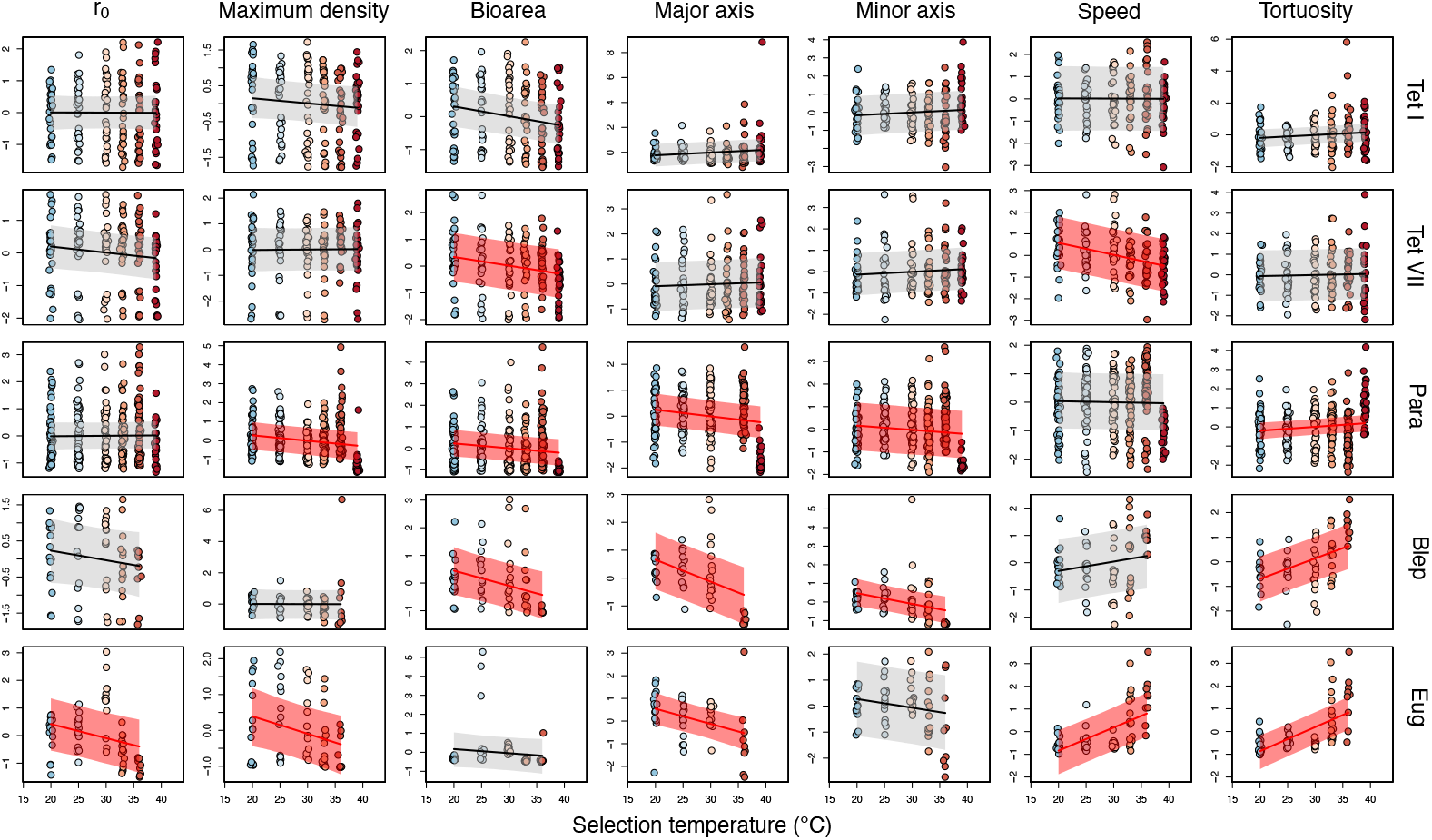
Median model predictions (solid lines) and 95% compatibility interval (shaded areas) obtained from the multivariate analysis of the 7 standardized phenotypic traits (*r*_0_, maximum density, bioarea, maximum axis, minimum axis, swimming speed and tortuosity) assayed at different temperatures ranging from 10 to 40°C. Each point is a standardized trait, and correspond to a biological replicate. The colours are the different selection temperature (evolutionary treatment of origin) as also indicated on the x-axis. Model prediction with parameter slopes different from 0 are highlighted in red. Tet I and VII are two mating types of *Tetrahymena thermophila*, Para is *Paramecium caudatum*, Blep is *Blepharisma* sp., and Eug is *Euglena gracilis*.

## Discussion

Climate change has had wide-ranging impacts on species, communities and ecosystems (Walther, 2010) and remains one of the biggest threats to biodiversity worldwide (Urban *et al*., 2016). Globally warming temperatures are well documented (Masson-Delmotte *et al*., 2021), making it important to understand if and how species can adapt to increased temperatures. This understanding is central for predicting the impacts of climate change on species persistence, extinction rates and changes to community compositions, for example.

In order to increase our understanding of evolutionary responses to temperature increases, we here experimentally evolved four different protist species at multiple increased temperatures and compared their thermal performance curves for different traits after one year of selection. In general, we found only weak signals for the evolution of population growth, but stronger responses for body size and movement.

### Evolution of population growth rate: weak signatures of adaptation

Using a Sharpe-Schoolfield model to fit TPCs on measured population growth rate data we only found weak signals of TPC evolution. This lack of a clear evolutionary response was unexpected, since all the protist species underwent selection for a year, in a selection regime far more intense than would be experienced in wild populations under the worst case scenarios for global warming (Masson-Delmotte *et al*., 2021). We did find signals of TPC evolution for growth rate in *T. thermophila*, more specifically 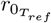 decreased with increasing selection temperature. We also report a general shift in niche parameter space and niche shape at the most extreme selection temperature (39°C; Fig. 1, Fig. S2).

We had particularly expected responses in *T. thermophila* because the (effective) population sizes in the selection lines were rather large (at equilibrium density approx 3.2 *×* 10^6^ cells at 20°C and 1.7 *×* 10^6^ at 39°C) and were selected for at least 857 generations. Furthermore, *T. thermophila* has an estimated macro-nuclear mutation rate (*U* = 0.0333 per haploid macronucleus genome per generation) that is orders of magnitudes higher than estimates for other protists, and eukaryotes in general (Brito *et al*., 2010). Therefore, we assumed that there should have been ample opportunity for evolution to occur. Finally, *T. thermophila* has been shown to successfully adapt to other selection pressures (e.g., pH Moerman *et al*. 2020a; and dispersal: Fronhofer *et al*. 2017) and has shown evolutionary change in response to temperature in previous studies (de Melo *et al*., 2020).

Most strikingly, *T*_*opt*_ and *r*_*max*_ were completely unaffected by our temperature treatments in all five species. Our results also do not support the “hotter-is-better” hypothesis, similarly to what was found in a meta-analysis on TPC evolution (Malusare *et al*., 2023). Several experimental studies in ectotherms have illustrated the rapid evolution of *T*_*opt*_ (Santos *et al*., 2006; Carbonell & Stoks, 2020; Mesas *et al*., 2021; Wang *et al*., 2023), but the associated evolution of *r*_*max*_ is usually inconsistent, and an overall clear consensus for evolutionary trajectories of TPCs is currently lacking. A recent evolutionary experiment in which eight species (51 genotypes) of the *Saccharomyces* genus were exposed to increasing temperatures for 600 generations found strong between- and within-species differences in TPC evolution. Interestingly, the authors identified two different evolutionary trajectories: genotypes either exhibited a “hotter is wider” or a “jack of all temperatures is a master of none” pattern (Molinet & Stelkens, 2025). This study highlights the complexity of adaptive responses to thermal selection with the presence of genotype-by-environmental interactions.

The absence of responses in our study and others may be explained by the biochemical adaptation (Montagnes *et al*., 2022) or the perfect compensation hypothesis (Frazier *et al*., 2006). These postulate that a fixed *r*_*max*_ may be the evolutionary result of compensating for thermodynamic constraints at a given temperature. The apparent absence of clearer responses could be also due to our current limitations in mechanistic theory, and therefore nonsensical estimates in TPC parameters. For instance, metabolic theory of ecology (Brown *et al*., 2004) has not yet integrated how the biochemical kinetics of the metabolism contribute to a single activation energy parameter *E* (O’Connor *et al*., 2007).

The lack of adaptation in growth rate to increased temperature in this study may suggest that adaptation to thermal stress from de novo mutations is more difficult than adaptation to other stressors, such as to pH (Moerman *et al*., 2020a), for example. While mechanistic targets of selection are less well defined for temperature, often observed strategies of adaptation to temperature include the evolution of ‘flexible’ enzymes (Marx *et al*., 2006), paralogous isozymes through duplication and mutation of genes (Somero, 1995), membrane composition (Hazel, 1995) and mitochondrial density/ capacity (Pörtner, 2001), for example. Importantly, as temperature affects the folding of proteins and the kinetics of biochemical reactions within a cells, it seems likely that a wide range of adaptations would be needed to confer thermal adaptation. Indeed, for example in *E. coli*, adaptation to extreme temperature was characterised by many epistatic interactions across the genome (Tenaillon *et al*., 2012). Further, a study of the thermal limits of 2000 taxa found that ancestral adaptation to cold temperatures happens far quicker than adaptation to warm temperatures (Bennett *et al*., 2021).

### Evolution of body size and swimming behaviour

The three largest protist species, *P. caudatum, Blepharisma* sp. and *E. gracilis*, exhibited consistent evolutionary changes in traits other than growth rate: Selection at the highest temperatures led to a reduction in their body size and modifications in swimming behaviour (swimming speed and tortuosity) across the assayed temperatures (Fig. 4). Changes in swimming speed were also found in one strain of *T. thermophila*.

We observed an overall reduction in body size of the evolved lines compared to ancestral conditions in all species (Fig. S4). Body size reduction might be adaptive and in line with expectations from Bergmann’s rule (Watt *et al*., 2010). This considers ecological and evolutionary responses of body size to temperatures, and hypothesizes increase size in colder climates as an evolutionary strategy to conserve heat due to lower surface-to-volume ratio in larger organisms, although this might not be universal (Angilletta & Dunham, 2003; Kingsolver & Huey, 2008). At warmer temperatures instead, smaller body sizes might be favoured because of their lower metabolic demands and efficiency in facing oxygen limitations (Gardner *et al*., 2011; Verberk *et al*., 2020). Thus, the reduction in body size in life-history diverse protist species might be a general response to the warmest temperatures experienced during the 1-year selection experiment. *T. thermophila* was the exception here, and this was in contrast to a previous study showing evolutionary decreases in body size in response to thermal selection (de Melo *et al*., 2020)). *T. thermophila* is the smallest of the four species in the study, and it may have been harder or even impossible to respond towards such evolutionary trajectory. It is possible that *T. thermophila* has already attained its biophysical limit due to laboratory handling, and could therefore not additionally decrease its body size. Several thermal experimental evolution studies have shown a direct relationship between increased temperature and the evolution of reduced sizes (Anderson, 1973; Cavicchi *et al*., 1989; Partridge *et al*., 1994), and this is also generally expected in aquatic ecosystems due to global warming (Daufresne *et al*., 2009). However, the general trend is still debated, and several additional factors should be considered such as intra- and inter-specific variation or the association with other traits, making predictions on body size evolution not straightforward. For instance, in their meta-analysis (Siepielski *et al*., 2019) found little support for adaptive evolutionary shift to shrinking body size in function of increased global temperatures.

The concurrent evolution of the swimming movement TPC is a result that should be interpreted carefully, especially considering how the experimental design had no explicitly spatial component. Previous evolutionary experiments with *P. caudatum* (Zilio *et al*., 2023b,a) and *T. thermophila* (Fronhofer & Altermatt, 2015) have provided evidence for the rapid evolution of swimming behaviour. Similarly to our findings showing for *P. caudatum* the evolution of higher tortuosity with warming temperatures, selection at the front of laboratory-simulated range expansions led to the evolution of higher dispersal, which was associated with the evolution of increased tortuosity (Zilio *et al*., 2023b,a). Tortuosity is the propensity to change directions while swimming, and can be interpreted as a more exploratory swimming behaviour, which could indeed facilitate dispersal or allow to closely track micro-variation in the environmental conditions. It is tempting to (i) speculate how selection at the highest temperature, likely representing the most stressful conditions, might have selected *P. caudatum* for increased tortuosity and potentially dispersal to more benign conditions, and (ii) to generalize this to *Blepharisma* sp. and *E. gracilis* which also showed the same pattern for reduced body size. However, in the studies mentioned above the trait under selection was dispersal, and evolutionary changes in swimming behaviour might be simply correlative responses or part of an emerging dispersal syndromes (Cote *et al*., 2016). Such speculative arguments do not necessarily hold for Tet VII. In *T. thermophila*, it is higher swimming speed rather than tortuosity to be associated with dispersal (Fronhofer & Altermatt, 2015), but our results suggest the evolution of reduced swimming speed at increased selection temperature. Nonetheless, *T. thermophila* already responded to laboratory conditions by increasing its swimming speed (Fig. S4). Thus, similarly to what was proposed before for body size, *T. thermophila* might have already reached some intrinsic limits and could not further increase swimming speed.

### Experimental design and limitations

Although we found evolutionary responses to thermal selection, it is important to put these results into context with some limits placed upon evolution in our experiment. Each replicate line was established from a clonal line without any standing genetic variation, a scenario that is unlikely to be the norm during adaptation to increased temperatures in wild populations. Standing genetic variation is expected to increase the speed of adaptation as i) the pace of evolution is a function of the amount of variation present, ii) the variation already present may be ‘pre-tested’ (i.e., it was not so deleterious as to be purged from the gene pool) and iii) there is a higher probability of fixation of weakly beneficial alleles as fixation probability is not only a function of the magnitude of the beneficial effect, but also of the effective population size (Barrett & Schluter, 2008). Further, as only a single mating type was present in each selection line, no recombination could occur, which many ciliates do under stressful conditions (Collins, 2012), and could be expected to increase the speed of adaptation (Cooper, 2007; McDonald *et al*., 2016). Our selection regime also imposed (comparatively mild) regular bottlenecks (1:10) during transfers to new bacterized medium, an often unavoidable feature of experimental evolution using serial transfer batch cultures (Wahl *et al*., 2002), which would act to limit the effective population size of the populations and therefore increase the effect of drift, reducing the probability that a beneficial mutation will reach fixation (Frankham *et al*., 1999).

Wild populations are unlikely to experience such strong, persistent selection pressures and constant temperatures, but rather fluctuations (e.g., due to diurnal and seasonal temperature cycles) or heatwaves (Mazdiyasni & AghaKouchak, 2015), which impact how we might expect populations to respond to climate change. Temperature fluctuations can have important consequences for biological processes (Slein *et al*., 2023). They can interact with other stressors (Verheyen *et al*., 2019; Chang *et al*., 2022) and modify biotic interactions (Raffel *et al*., 2012), leading to different outcomes than constant thermal regimes (for a discussion for adaptation to fluctuating temperatures see also Malusare *et al*., 2023). In a recent meta-analysis and synthesis, Stocker *et al*. (2024) found that biological responses to thermal change were mostly driven by mean temperatures and not fluctuations, but this was context-dependent. For instance, in freshwater environments (protist habitat), the general pattern was indeed reversed, and the fluctuating temperatures had a stronger impact on biological responses than stable temperatures, suggesting how fluctuations might be the most relevant temperature regime to be investigated using protists.

### Implications and conclusions

While both local adaptation and dispersal (movement) in response to a warming world will undoubtedly play a role in species’ persistence (see e.g., Norberg *et al*., 2012; Thompson & Fronhofer, 2019; Usui & Angert, 2024), there are many cases in which dispersal may be too limited, for instance due to anthropogenic fragmentation and loss of connecting habitat (Harrison & Bruna, 1999; Fahrig, 2017; Fletcher *et al*., 2018; Kamal *et al*., 2025). In such cases, understanding how a population can be rescued (Gonzalez *et al*., 2013; Chevin & Bridle, 2025) from environmentally induced declines via adaptation is vital to truly evaluate the potential effects of climate change. Reduction in body size, for example, could be adaptive against extreme temperature, but it may have detrimental consequences, affecting eco-evolutionary dynamics and ecosystem functioning (Dossena *et al*., 2012). Populations with reduced body size may be more exposed to other global change stressors (e.g., pollutants) or suffer more from antagonistic biotic interactions (e.g., parasites or predators), potentially increasing their extinction probability. In fact, change towards smaller body sizes is even used as a proxy and predictor to detect and manage populations at risk of collapse (Cardillo, 2021; Williams *et al*., 2021).

In this study, we have assayed the population growth of multiple protist species that have undergone selection at increased temperatures. Our results highlight the importance of using multiple-traits approaches to investigate a species response to increasing temperatures and TPC evolution, and have implications for how species may respond to climate change and impact whole ecosystems.

## Author contributions

E.A.F. and S.P.M. conceived the study. S.P.M., I.R.M., M.-A.D., J.G. and C.G.-B. gathered the data. G.Z. and I.R.M. performed the statistical analyses. G. Z., I.R.M. and E.A.F. wrote the manuscript and all authors commented on the draft.

## Acknowledgements

This work was supported by a grant from the Agence Nationale de la Recherche (No.: ANR-19-CE02-0015) to EAF. This is publication ISEM-YYYY-XXX of the Institut des Sciences de l’Evolution – Montpellier.

## Code and Data availability

Data and R-code will be made available upon publication.

## Notes

### Competing Interest Statement

The authors have declared no competing interest.

