## Supplementary Material for "Experimental Evolution of the Thermal Performance Curve"

### S1 Supplementary methods

#### S1.1 R code

```r
# Script for fitting Sharpe-Schoolfield niche functions with stan via brms.
# Priors changed depending on the species
# clear environment ---------------------------------------------------------
rm(list=ls())
# load packages --------------------------------------------------------------
library(tidyverse)
library(rstan)
library(janitor)
library(patchwork)
library(brms)
library(tidybayes)
# options --------------------------------------------------------------------
## stan options
rstan_options(auto_write = TRUE)
options(mc.cores = parallel::detectCores())
# Import dataset from set directory
  ----------------------------------------------------------------------------
data<-read.table(file="file.txt")
# Experimental details changed depending on status, species, replicate, selection
#temperature, and year
d<-data[data$Evo_Status=="Evolved" &
data$species2=="Blep_I" &
data$replicate == "c" &
data$selection_temperature == "25",]
d$tK <- (d$temperature) + 273.15
d$tref <- rep(20 + 273.15,length(d$selection_temperature))
d$k <- rep(8.62e-05,length(d$selection_temperature))
nlform <- bf(r0_median|se(r0_sd, sigma = TRUE) ~ (exp(rtref) * exp(exp(e)/k * (1/tref -
1/(tK)))) * (1/(1 + exp(-exp(el)/k * (1/(tl+273.15) - 1/(tK)))+exp(exp(eh)/k * (1/(th
  +273.15) -
1/(tK))))),
rtref ~ 1,
e ~ 1,
el ~ 1,
tl ~ 1,
eh ~ 1,
th ~ 1,
nl = TRUE) nlprior <- c(prior(normal(log(0.2), 0.4), nlpar = "rtref"),
prior(normal(log(0.25), 0.6), nlpar = "e"),
```

```r
prior(normal(log(2.5), 0.75), nlpar = "el"),
prior(normal(15,4), nlpar = "tl"),
prior(normal(log(2.5), 0.75), nlpar = "eh"),
prior(normal(35,4), nlpar = "th"))
fit_test_mod <- brm(formula = nlform,
data = d,
family = gaussian(),
prior = nlprior,
control = list(adapt_delta = 0.99),
chains = 4,
iter = 10000
)
summary(fit_test_mod)
# Script for fitting niche functions with stan via brms
# clear environment ----------------------------------------------------------
rm(list=ls())
# load packages --------------------------------------------------------------
library(tidyverse)
library(rstan)
library(janitor)
library(patchwork)
library(brms)
library(tidybayes)
# options --------------------------------------------------------------------
## stan options
rstan_options(auto_write = TRUE)
options(mc.cores = parallel::detectCores())
d <- read_csv("file.csv") %>%
group_by(speciesMT, replicate, selection_temperature) %>%
mutate(n = n()) %>%
filter(n >= 7) %>%
ungroup()d <- d %>%
filter(speciesMT == "Tet_VII", replicate == "a", selection_temperature == 39) %>%
mutate(tK = temperature + 273.15,
tref = 20 + 273.15,
k = 8.62e-05) %>%
select(speciesMT, replicate, selection_temperature, temperature, tK, tref, k, r0_median
 ,
r0_sd)
nlform <- bf(r0_median|se(r0_sd, sigma = TRUE) ~ (exp(rtref) * exp(exp(e)/k * (1/tref -
1/(tK)))) * (1/(1 + exp(-exp(el)/k * (1/(tl+273.15) - 1/(tK)))+exp(exp(eh)/k * (1/(th
  +273.15) -
```

```
1/(tK))))),
rtref ~ 1,
e ~ 1,
el ~ 1,
tl ~ 1,
eh ~ 1,
th ~ 1,
nl = TRUE)
nlprior <- c(prior(normal(log(0.2), 0.4), nlpar = "rtref"),
prior(normal(log(0.25), 0.6), nlpar = "e"),
prior(normal(log(2.5), 0.75), nlpar = "el"),
prior(normal(15,4), nlpar = "tl"),
prior(normal(log(2.5), 0.75), nlpar = "eh"),
prior(normal(35,4), nlpar = "th"))
fit_test_mod <- brm(formula = nlform,
data = d,
family = gaussian(),
prior = nlprior,
control = list(adapt_delta = 0.99),
chains = 4,
iter = 10000
)
summary(fit_test_mod)
prior_dist <-
exp(rnorm(1:80000,log(0.2),0.4)) %>%
bind_cols() %>%
rename("b_rtref_Intercept" = "...1") %>%
bind_cols()
prior_dist <-
exp(rnorm(1:80000,log(0.25), 0.6)) %>%
bind_cols() %>%
rename("b_e_Intercept" = "...1") %>%bind_cols(prior_dist)
prior_dist <-
exp(rnorm(1:80000,log(2.5), 0.75)) %>%
bind_cols() %>%
rename("b_el_Intercept" = "...1") %>%
bind_cols(prior_dist)
prior_dist <-
rnorm(1:80000,15,3) %>%
bind_cols() %>%
# filter(...1 < 25) %>%
rename("b_tl_Intercept" = "...1") %>%
```

```
bind_cols(prior_dist)
prior_dist <-
exp(rnorm(1:80000,log(2.5), 0.75)) %>%
bind_cols() %>%
rename("b_eh_Intercept" = "...1") %>%
bind_cols(prior_dist)
prior_dist <-
rnorm(1:80000,35,4) %>%
bind_cols() %>%
# filter(...1 > 25) %>%
rename("b_th_Intercept" = "...1") %>%
bind_cols(prior_dist) %>%
gather(., 'param', 'estimate', 1:ncol(.)) %>%
mutate(type = "prior")
dists <- fit_test_mod %>%
spread_draws(`b_.*`, regex = TRUE) %>%
gather(., 'param', 'estimate', 4:ncol(.)) %>%
mutate(type = "post") %>%
bind_rows(prior_dist)
ggplot(dists, aes(x = estimate)) +
facet_wrap(~param, scales = "free") +
geom_density(aes(colour = type))
params <- fit_test_mod %>%
spread_draws(`b_.*`, regex = TRUE) %>%
gather(., 'param', 'estimate', 4:ncol(.)) %>%
separate(., param, c('blah', 'term', 'blah2'), sep = '_') %>%
select(., -starts_with('blah')) %>%
filter(., !is.nan(estimate)) %>%
group_by(., term) %>%mean_qi()
preds <- data.frame(tK = seq(0 + 273.15, 50 + 273.15, length.out = 200),
tref = 20 + 273.15,
k = 8.62e-05,
r0_sd = 1) %>%
add_fitted_draws(fit_test_mod, re_formula = NA) %>%
data.frame() %>%
group_by(tK) %>%
mean_qi(estimate = .value)
ggplot(preds, aes(x = tK-273.15, y = estimate)) +
geom_line() +
geom_ribbon(aes(ymin = .lower, ymax = .upper), alpha = 0.2) +
geom_pointrange(data = d, aes(x = temperature, y = r0_median, ymin = r0_median-r0_sd,
ymax = r0_median+r0_sd))
```

```r
stancode(fit_test_mod)
plot(fit_test_mod)
pairs(fit_test_mod)
fit_test_params <- fit_test_mod %>%
spread_draws(`b_.*`, regex = TRUE)
temps <- seq(15,50,0.1)+273.15
values <- tibble()
head(fit_test_params)
for (temp in temps) {
  print(temp)
  t <-
  fit_test_params %>%mutate(topt_temp = temp,
  value = (b_rtref_Intercept * exp(b_e_Intercept/8.62e-05 * (1/20 - 1/(temp)))) * (1/(1
   +
  exp(-b_el_Intercept/8.62e-05 * (1/(b_tl_Intercept+273.15) -
  1/(temp)))+exp(b_eh_Intercept/8.62e-05 * (1/(b_th_Intercept+273.15) - 1/(temp))))))
  values <- bind_rows(values, t)
}
topt_range <- values %>%
na_if(Inf) %>%
group_by(.chain,.iteration,.draw, b_rtref_Intercept, b_el_Intercept, b_tl_Intercept,
b_eh_Intercept, b_th_Intercept) %>%
filter(value == max(value)) %>%
distinct(.chain,.iteration,.draw, b_rtref_Intercept, b_el_Intercept, b_tl_Intercept,
b_eh_Intercept, b_th_Intercept, .keep_all = TRUE) %>%
rename("b_topt_Intercept" = "topt_temp",
"b_rmax_Intercept" = "value") %>%
gather(., 'param', 'estimate', 4:ncol(.)) %>%
separate(., param, c('blah', 'term', 'blah2'), sep = '_') %>%
select(., -starts_with('blah')) %>%
filter(., !is.nan(estimate)) %>%
group_by(., term) %>%
mean_qi()
preds <- data.frame(tK = seq(5 + 273.15, 40 + 273.15, length.out = 200),
tref = 20 + 273.15,
k = 8.62e-05,
r0_sd = 1) %>%
add_fitted_draws(fit_test_mod, re_formula = NA) %>%
data.frame() %>%
group_by(tK) %>%
mean_qi(estimate = .value)
Topt <- topt_range %>%
```

```
filter(term == "topt") %>%
rename("est" = "estimate")
ggplot(preds, aes(x = tK-273.15, y = estimate)) +
geom_line() +
geom_ribbon(aes(ymin = .lower, ymax = .upper), alpha = 0.2) +
geom_pointrange(data = d, aes(x = temperature, y = r0_median, ymin = r0_median-r0_sd,
ymax = r0_median+r0_sd)) +
geom_vline(data = Topt, aes(xintercept = est -273.15)) +
geom_vline(data = Topt, aes(xintercept = .upper -273.15)) +
geom_vline(data = Topt, aes(xintercept = .lower -273.15))
r_tref * exp(e/k * (1/tref - 1/tK))*1/(1 + exp(eh/k * (1/(th + 273.15) - 1/tK)))nlform
  <- bf(r0_median|se(r0_sd, sigma = TRUE) ~ (rtref * exp(e/k * (1/tref - 1/tK))*1/(1 +
exp(eh/k * (1/(th + 273.15) - 1/tK)))),
rtref ~ 1,
e ~ 1,
eh ~ 1,
th ~ 1,
nl = TRUE)
nlprior <- c(prior(normal(0.2,5), nlpar = "rtref", lb = 0),
prior(normal(0.005,5), nlpar = "e", lb = 0),
prior(normal(6.5,5), nlpar = "eh", lb = 0),
prior(normal(37,5), nlpar = "th", lb = 0))
fit_test_mod_2 <- brm(formula = nlform,
data = d,
prior = nlprior,
control = list(adapt_delta = 0.99),
chains = 4,
iter = 20000
)
summary(fit_test_mod_2)
stancode(fit_test_mod_2)
plot(fit_test_mod_2)
fit_test_params_2 <- fit_test_mod_2 %>%
spread_draws(`b_.*`, regex = TRUE) %>%
mutate(., b_Topt_Intercept = get_topt(b_eh_Intercept, b_th_Intercept, b_e_Intercept))
%>%
gather(., 'param', 'estimate', 4:ncol(.)) %>%
separate(., param, c('blah', 'term', 'blah2'), sep = '_') %>%
select(., -starts_with('blah')) %>%
filter(., !is.nan(estimate)) %>%
group_by(., term) %>%
mean_qi()
```

```
preds_2 <- data.frame(tK = seq(5 + 273.15, 40 + 273.15, length.out = 200),
tref = 20 + 273.15,
k = 8.62e-05,
r0_sd = 1) %>%
add_fitted_draws(fit_test_mod_2, re_formula = NA) %>%
data.frame() %>%
group_by(tK) %>%
mean_qi(estimate = .value)Topt_2 <- fit_test_params_2 %>%
filter(term == "Topt") %>%
rename("est" = "estimate")
get_topt <- function(Eh, Th, Ea){
  return((Eh * Th)/(Eh + (8.62e-05 * Th * log((Eh/Ea) - 1))))
}
get_topt_2 <- function(rtref, e-
(605635882065044123207854847514308328942767351791616 * rtref * e^(-31534.6/tK) *
((eh - 2.71828) * e^(11600.9 * eh * (1/th - 1/tK)) - 2.71828)) / (tK^2 (e^(11600.9 * eh
   * (1/th
- 1/tK)) + 1)^2)
get_topt(4.3707442, 32.1593038, 0.8663518)
ggplot(preds_2, aes(x = tK-273.15, y = estimate)) +
geom_line(aes(x = tK-273.15, y = estimate)) +
geom_ribbon(aes(ymin = .lower, ymax = .upper), alpha = 0.2) +
geom_vline(aes(xintercept = 32.13))
geom_line(data = preds_2, aes(x = tK-273.15, y = estimate), colour = "blue") +
geom_ribbon(data = preds_2, aes(ymin = .lower, ymax = .upper), alpha = 0.2, fill = "
  blue") +
geom_pointrange(data = d, aes(x = temperature, y = r0_median, ymin = r0_median-r0_sd,
ymax = r0_median+r0_sd)) +
geom_vline(data = Topt_2, aes(xintercept = est), colour = "blue") +
geom_vline(data = Topt_2, aes(xintercept = .upper), colour = "blue", linetype = "dashed
  ") +
geom_vline(data = Topt_2, aes(xintercept = .lower), colour = "blue", linetype = "dashed
  ") +
geom_vline(data = Topt, aes(xintercept = est), colour = "grey") +
geom_vline(data = Topt, aes(xintercept = .upper), colour = "grey", linetype = "dashed")
   +
geom_vline(data = Topt, aes(xintercept = .lower), colour = "grey", linetype = "dashed")
geom_ribbon(aes(ymin = .lower, ymax = .upper), alpha = 0.2) +
geom_pointrange(data = d, aes(x = temperature, y = r0_median, ymin = r0_median-r0_sd,
ymax = r0_median+r0_sd)) +
geom_vline(data = Topt, aes(xintercept = est)) +
geom_vline(data = Topt, aes(xintercept = .upper)) +
```

```
geom_vline(data = Topt, aes(xintercept = .lower))
```

### S2 Supplementary figures

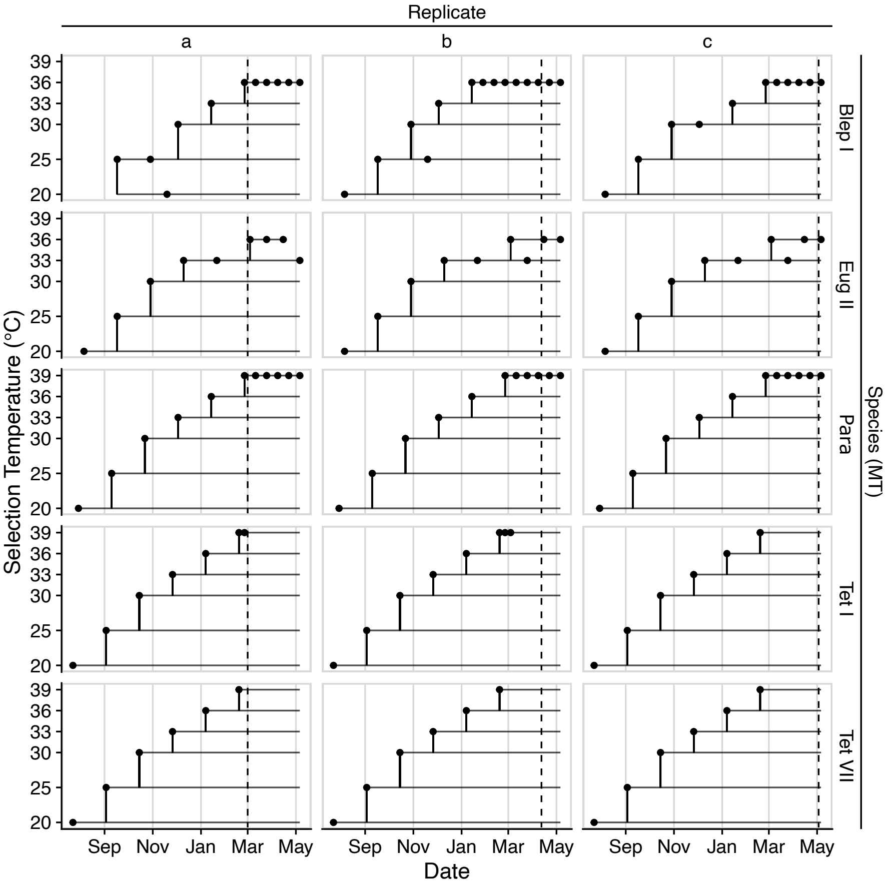


Figure S1: Chronogram showing the selection experiment history. Vertical lines between selection temperatures represent an initial transfer. Dots represent extinctions where the population had to be re-established from the previous selection temperature. Dashed lines show the starting assay date for each population.

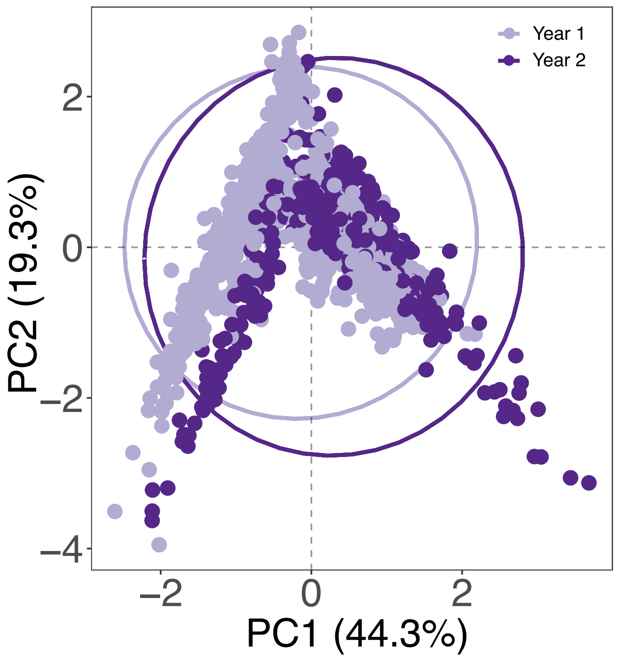


Figure S2: PCA with 95% confidence ellipsis probability of the 7 phenotypic traits for all species and selection temperatures (complete dataset). The two colours show the first and second year of selection.

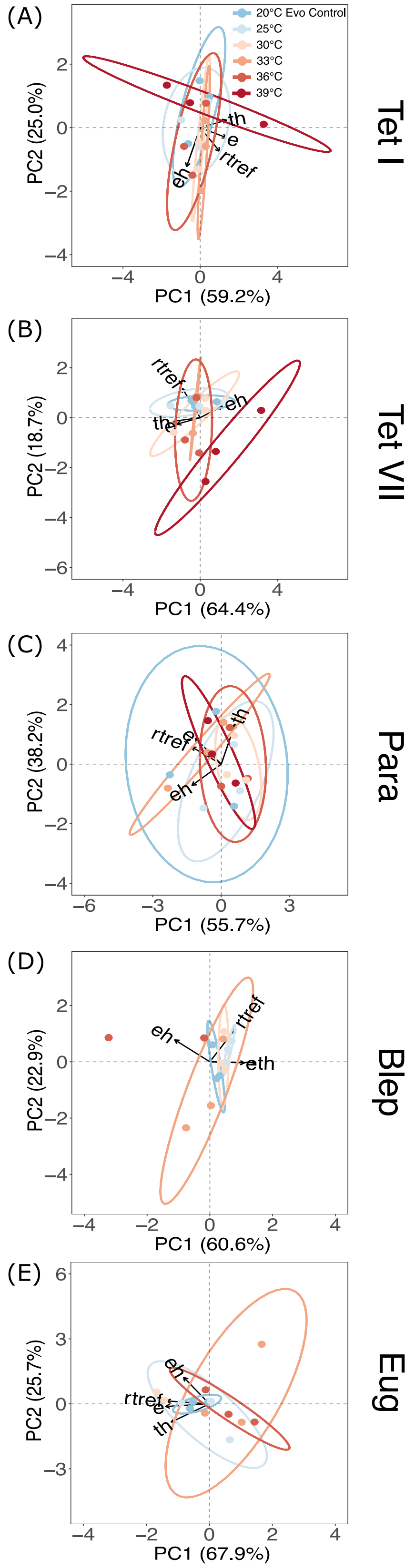


Figure S3: PCA with 95% confidence ellipsis probability of the TPC parameters. Colours are the selection temperature regime. Each point represents the average parameter value in multi-variate space of a biological replicate, with arrows indicating the direction of trait variation. Tet I and VII are two mating types of *Tetrahymena thermophila*, Para is *Paramecium caudatum*, Blep is *Blepharisma* sp., and Eug is *Euglena gracilis*.

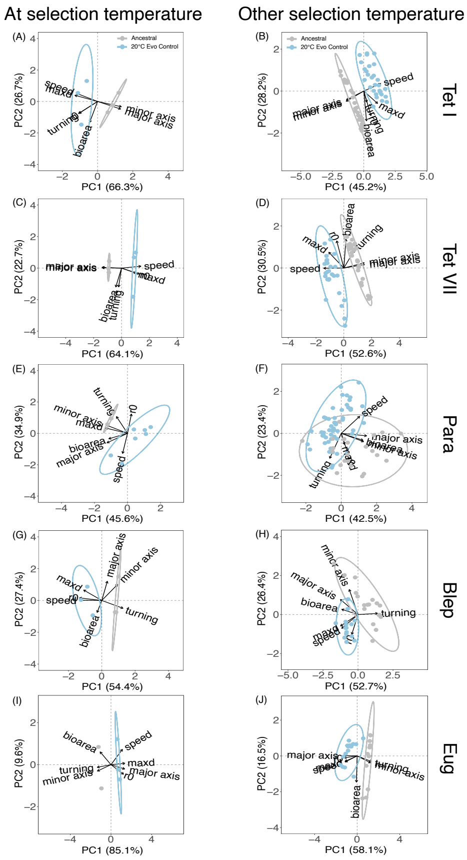


Figure S4: PCA with 95% confidence ellipsis probability of the 7 phenotypic traits measured at 20°C (A, C, E, G, I) or at other temperatures (B, D, F, H, J) for the ancestral (grey) and evolutionary control lines (light blue). Each of points represents the average phenotypic value in multi-variate space, with arrows indicating the direction of trait variation. Tet I and VII are two mating types of *Tetrahymena thermophila*, Para is *Paramecium caudatum*, Blep is *Blepharisma* sp., and Eug is *Euglena gracilis*.

### S3 Supplementary tables

Table S1: Number of selection lines over the two-year experiment. Extinction events and total number of lines available at the end of each year per species and selection temperature are in bold.

| **Species** | **Year 1** | | | | | | | **Year 2** | | | | | | |
|---|---|---|---|---|---|---|---|---|---|---|---|---|---|---|
| | 20°C | 25°C | 30°C | 33°C | 36°C | 39°C | Tot | 20°C | 25°C | 30°C | 33°C | 36°C | 39°C | Final |
| Tet I | 3/3 | 3/3 | 3/3 | 3/3 | 3/3 | 3/3 | 18/18 | 3/3 | 3/3 | 3/3 | 3/3 | 3/3 | 3/3 | 18/18 |
| Tet VII | 3/3 | 3/3 | 3/3 | 3/3 | 3/3 | 3/3 | 18/18 | 3/3 | 3/3 | 3/3 | 3/3 | 3/3 | **2/3** | **17/18** |
| Para | 3/3 | 3/3 | 3/3 | 3/3 | 3/3 | 3/3 | 18/18 | 3/3 | 3/3 | 3/3 | 3/3 | 2/3 | **0/3** | **14/18** |
| Blep | 3/3 | 3/3 | 3/3 | 3/3 | **2/3** | **0/3** | **14/18** | 3/3 | 3/3 | 3/3 | **2/3** | **0/3** | **0/3** | **11/18** |
| Eug | 3/3 | 3/3 | 3/3 | 3/3 | 3/3 | **0/3** | **15/18** | 3/3 | 3/3 | 3/3 | **2/3** | **1/3** | **0/3** | **12/18** |

Table S2: Priors for fitting the Sharpe-Schoolfield thermal reaction norm in Eq. 1 of the main text.

| Species | e | $e_h$ | $t_h$ | $r_{ref}$ |
|---|---|---|---|---|
| Tet I | normal(log(0.25), 0.6) | normal(log(2.5), 0.75) | normal(35,4) | normal(log(0.2), 0.4) |
| Tet VII | normal(log(0.25), 0.6) | normal(log(2.5), 0.75) | normal(35,4) | normal(log(0.2), 0.4) |
| Para | normal(log(0.25), 0.6) | normal(log(2.5), 0.75) | normal(35,4) | normal(log(0.025), 0.4) |
| Blep | normal(log(0.25), 0.6) | normal(log(2.5), 0.75) | normal(35,4) | normal(log(0.025), 0.4) |
| Eug | normal(log(0.25), 0.6) | normal(log(2.5), 0.75) | normal(35,4) | normal(log(0.025), 0.4) |

Table S3: Statistical models for the multi-variate analysis of niche parameters comparing ancestral and evolutionary control (20°C). Each row is a different model; the best model is highlighted in bold.

| Species | Model | Selection line random term | Error propagation | WAIC | Δ WAIC | SE | WAIC weight |
|---|---|---|---|---|---|---|---|
| Tet I | Intercept | No | No | 75.44 | 8.03 | 6.92 | 0.02 |
| | **Evolutionary conditions** | **No** | **No** | **67.41** | | **6.39** | **0.98** |
| Tet VII | Intercept | No | No | 71.16 | 57.87 | 6.84 | 0 |
| | **Evolutionary conditions** | **No** | **No** | **57.87** | | **6.98** | **1** |
| Para | Intercept | No | No | 65.96 | 1.59 | 7.00 | 0.31 |
| | **Evolutionary conditions** | **No** | **No** | **64.37** | | **6.61** | **0.69** |
| Blep | Intercept | No | No | 63.35 | 11.84 | 3.10 | 0 |
| | **Evolutionary conditions** | **No** | **No** | **51.51** | | **3.31** | **1** |
| Eug | Intercept | No | No | 45.90 | 6.13 | 4.76 | 0.04 |
| | **Evolutionary conditions** | **No** | **No** | **39.77** | | **3.41** | **0.96** |

Table S4: Statistical models for the multi-variate analysis of niche parameters comparing lines evolved at different selection temperatures (including evolutionary control). Each row is a different model; the best model is in bold.

| Species | Model | Selection line random term | Error propagation | WAIC | Δ WAIC | SE | WAIC weight |
|---|---|---|---|---|---|---|---|
| Tet I | Intercept | Yes | No | 198.64 | 5.77 | 22.95 | 0.05 |
| | **Selection temperature** | **Yes** | **No** | **192.87** | | **21.45** | **0.95** |
| Tet VII | Intercept | Yes | No | 180.4 | 7.7 | 23.69 | 0.02 |
| | **Selection temperature** | **Yes** | **No** | **172.7** | | **19.89** | **0.98** |
| Para | **Intercept** | **Yes** | **No** | **169.28** | 2.81 | **15.25** | **0.8** |
| | Selection temperature | Yes | No | 172.09 | | 15.02 | 0.2 |
| Blep | **Intercept** | **Yes** | **No** | **135.49** | 4.03 | **20.82** | **0.88** |
| | Selection temperature | Yes | No | 139.52 | | 17.50 | 0.12 |
| Eug | **Intercept** | **Yes** | **No** | **117.15** | 4.81 | **15.57** | **0.92** |
| | Selection temperature | Yes | No | 121.96 | | 13.43 | 0.08 |

Table S5: Median slope and 95% compatibility intervals obtained from posterior distributions of the multi-variate analysis of niche parameters comparing lines evolved at different selection temperatures (including evolutionary control). Slopes different from 0 are highlighted in bold.

| Species | $e$ | $e_h$ | $t_h$ | $r_{ref}$ |
|---|---|---|---|---|
| Tet I | 0.048<br>(-0.020; 0.118) | 0.036<br>(-0.044; 0.116) | 0.024<br>(-0.052; 0.102) | -0.027<br>(-0.113; 0.056) |
| Tet VII | -0.017<br>(-0.082; 0.047) | 0.006<br>(-0.074; 0.086) | -0.049<br>(-0.120; 0.021) | **-0.096**<br>**(-0.166; -0.027)** |
| Para | -0.012<br>(-0.075; 0.050) | -0.026<br>(-0.090; 0.037) | 0.040<br>(-0.018; 0.100) | 0.008<br>(-0.069; 0.084) |
| Blep | -0.081<br>(-0.188; 0.026) | 0.049<br>(-0.058; 0.157) | -0.082<br>(-0.172; 0.006) | -0.034<br>(-0.150; 0.079) |
| Eug | -0.052<br>(-0.148; 0.042) | -0.002<br>(-0.130; 0.090) | -0.065<br>(-0.159; 0.027) | -0.068<br>(-0.157; 0.021) |

Table S6: Statistical models for the analysis of $T_{opt}$ comparing ancestral and evolutionary control (20°C). Each row is a different model; the best model is highlighted in bold.

| Species | Model | Selection line random term | Error propagation | WAIC | Δ WAIC | SE | WAIC weight |
|---|---|---|---|---|---|---|---|
| Tet I | **Intercept** | **No** | **Yes** | **27.06** | 1.44 | **0.29** | **0.67** |
| | Evolutionary conditions | No | Yes | 28.50 | | 0.27 | 0.33 |
| Tet VII | **Intercept** | **No** | **Yes** | **28.5** | 1.29 | **0.21** | **0.65** |
| | Evolutionary conditions | No | Yes | 29.79 | | 0.21 | 0.35 |
| Para | **Intercept** | **No** | **Yes** | **31.60** | 0.22 | **1,38** | **0.53** |
| | Evolutionary conditions | No | Yes | 31.82 | | 1.17 | 0.47 |
| Blep | **Intercept** | **No** | **Yes** | **28.76** | 1.28 | **0.14** | **0.65** |
| | Evolutionary conditions | No | Yes | 30.04 | | 0.12 | 0.35 |
| Eug | **Intercept** | **No** | **Yes** | **29.01** | 1.02 | **0.12** | **0.63** |
| | Evolutionary conditions | No | Yes | 30.03 | | 0.12 | 0.37 |

Table S7: Statistical models for the analysis of $T_{opt}$ comparing lines evolved at different selection temperatures (including evolutionary control). Each row is a different model; the best model is highlighted in bold.

| Species | Model | Selection line random term | Error propagation | WAIC | Δ WAIC | SE | WAIC weight |
|---|---|---|---|---|---|---|---|
| Tet I | **Intercept** | **Yes** | **Yes** | **77.74** | 1.22 | **1.13** | **0.65** |
| | Selection temperature | Yes | Yes | 78.96 | | 1.22 | 0.35 |
| Tet VII | **Intercept** | **Yes** | **Yes** | **83.05** | 1.18 | **0.38** | **0.64** |
| | Selection temperature | Yes | Yes | 84.23 | | 0.33 | 0.36 |
| Para | **Intercept** | **Yes** | **Yes** | **91.04** | 1.27 | **0.81** | **0.65** |
| | Selection temperature | Yes | Yes | 92.31 | | 1.02 | 0.35 |
| Blep | **Intercept** | **Yes** | **Yes** | **67.21** | 1.09 | **0.81** | **0.63** |
| | Selection temperature | Yes | Yes | 68.30 | | 0.75 | 0.37 |
| Eug | **Intercept** | **Yes** | **Yes** | **67.07** | 1.13 | **0.67** | **0.64** |
| | Selection temperature | Yes | Yes | 68.20 | | 0.59 | 0.36 |

Table S8: Statistical models for the analysis of $r_{max}$ comparing ancestral and evolutionary control (20°C). Each row is a different model; the best model is highlighted in bold.

| Species | Model | Selection line random term | Error propagation | WAIC | Δ WAIC | SE | WAIC weight |
|---|---|---|---|---|---|---|---|
| Tet I | **Intercept** | **No** | **Yes** | **0.19** | 1.94 | **0.45** | **0.72** |
| | Evolutionary conditions | No | Yes | 2.13 | | 0.42 | 0.28 |
| Tet VII | **Intercept** | **No** | **Yes** | **5.31** | 1.74 | **0.51** | **0.7** |
| | Evolutionary conditions | No | Yes | 7.05 | | 0.58 | 0.3 |
| Para | Intercept | No | Yes | -19.49 | 0.6 | 1.01 | 0.43 |
| | **Evolutionary conditions** | **No** | **Yes** | **-20.09** | | **1.18** | **0.57** |
| Blep | **Intercept** | **No** | **Yes** | **-17.14** | 1.64 | **1.19** | **0.69** |
| | Evolutionary conditions | No | Yes | -15.50 | | 1.19 | 0.31 |
| Eug | **Intercept** | **No** | **Yes** | **-14.28** | 1.42 | **1.10** | **0.67** |
| | Evolutionary conditions | No | Yes | -12.86 | | 1.11 | 0.33 |

Table S9: Statistical models for the analysis of $r_{max}$ comparing lines evolved at different selection temperatures (including evolutionary control). Each row is a different model; the best model is highlighted in bold.

| Species | Model | Selection line random term | Error propagation | WAIC | Δ WAIC | SE | WAIC weight |
|---|---|---|---|---|---|---|---|
| Tet I | **Intercept** | **Yes** | **Yes** | **-4.56** | 1.57 | **2.47** | **0.69** |
| | Selection temperature | Yes | Yes | -2.99 | | 2.38 | 0.31 |
| Tet VII | **Intercept** | **Yes** | **Yes** | **5.56** | 1.02 | **1.82** | **0.62** |
| | Selection temperature | Yes | Yes | 6.58 | | 1.55 | 0.38 |
| Para | **Intercept** | **Yes** | **Yes** | **-78.91** | 1.15 | **3.40** | **0.64** |
| | Selection temperature | Yes | Yes | -77.76 | | 3.27 | 0.36 |
| Blep | **Intercept** | **Yes** | **Yes** | **-35.20** | 0.23 | **0.87** | **0.6** |
| | Selection temperature | Yes | Yes | -35.43 | | 0.46 | 0.4 |
| Eug | **Intercept** | **Yes** | **Yes** | **-27.48** | 0.56 | **2.20** | **0.57** |
| | Selection temperature | Yes | Yes | -26.92 | | 2.27 | 0.43 |

Table S10: Statistical models for the multi-variate analysis of seven phenotypic traits. The analysis compares ancestral and evolutionary control (20°C) lines assayed at 20°C. Each row is a different model; the best model is highlighted in bold.

| Species | Model | Selection line random term | WAIC | Δ WAIC | SE | WAIC weight |
|---|---|---|---|---|---|---|
| Tet I | Intercept | No | 102.52 | 30.41 | 6.59 | 0 |
| | **Evolutionary conditions** | **No** | **72.11** | | **8.68** | **1** |
| Tet VII | Intercept | No | 99.01 | 63.11 | 7.71 | 0 |
| | **Evolutionary conditions** | **No** | **35.90** | | **9.61** | **1** |
| Para | Intercept | No | 121.22 | 3.11 | 4.62 | 0.17 |
| | **Evolutionary conditions** | **No** | **118.11** | | **4.42** | **0.83** |
| Blep | Intercept | No | 119.34 | 19.39 | 5.58 | 0 |
| | **Evolutionary conditions** | **No** | **99.95** | | **5.62** | **1** |
| Eug | Intercept | No | 87.72 | 37.29 | 5.50 | 0 |
| | **Evolutionary conditions** | **No** | **50.43** | | **3.03** | **1** |

Table S11: Statistical models for the multi-variate analysis of seven phenotypic traits. The analysis compares ancestral and evolutionary control (20°C) lines assayed at the other selection temperatures. Each row is a different model; the best model is highlighted in bold.

| Species | Model | Selection line random term | WAIC | Δ WAIC | SE | WAIC weight |
|---|---|---|---|---|---|---|
| Tet I | Intercept | No | 734.33 | 164.28 | 31.05 | 0 |
| | **Evolutionary conditions** | **No** | **570.05** | | **37.83** | **1** |
| Tet VII | Intercept | No | 665.65 | 183.07 | 32.69 | 0 |
| | **Evolutionary conditions** | **No** | **482.58** | | **39.31** | **1** |
| Para | Intercept | No | 1213.61 | 153.43 | 46.31 | 0 |
| | **Evolutionary conditions** | **No** | **1060.18** | | **53.78** | **1** |
| Blep | Intercept | No | 490.16 | 112.01 | 28.56 | 0 |
| | **Evolutionary conditions** | **No** | **378.15** | | **35.22** | **1** |
| Eug | Intercept | No | 434.57 | 95.27 | 22.29 | 0 |
| | **Evolutionary conditions** | **No** | **339.30** | | **30.78** | **1** |

Table S12: Statistical models for the multi-variate analysis of seven phenotypic traits. The analysis compares lines evolved at different selection temperatures (including evolutionary control) assayed at their own selection temperatures. Each row is a different model; the best model is highlighted in bold.

| Species | Model | Selection line random term | WAIC | Δ WAIC | SE | WAIC weight |
|---|---|---|---|---|---|---|
| Tet I | Intercept | Yes | 332.13 | 21.35 | 16.06 | 0 |
| | **Selection temperature** | **Yes** | **310.78** | | **14.72** | **1** |
| Tet VII | **Intercept** | **Yes** | **352.58** | 0.57 | **23.31** | **0.57** |
| | Selection temperature | Yes | 353.15 | | 21.13 | 0.43 |
| Para | **Intercept** | **Yes** | **262.72** | 4.09 | **11.59** | **0.89** |
| | Selection temperature | Yes | 266.81 | | 9.99 | 0.11 |
| Blep | Intercept | Yes | 249.85 | 10.16 | 7.84 | 0.01 |
| | **Selection temperature** | **Yes** | **239.69** | | **7.42** | **0.99** |
| Eug | **Intercept** | **Yes** | **245.27** | 6.41 | **14.17** | **0.96** |
| | Selection temperature | Yes | 251.68 | | 13.15 | 0.04 |

Table S13: Median slope and 95% compatibility intervals obtained from posterior distributions of the multi-variate analysis of 7 traits. The analysis compares lines evolved at different selection temperatures (including evolutionary control) assayed at their own selection temperatures. Slopes different from 0 are in bold.

| Species | $r_0$ | Max density | Bioarea | Major axis | Minor axis | Speed | Tortuosity |
|---|---|---|---|---|---|---|---|
| Tet I | 0.012<br>(-0.065; 0.098) | **-0.121**<br>**(-0.169; -0.072)** | -0.056<br>(-0.134; 0.022) | -0.038<br>(-0.118; 0.041) | -0.012<br>(-0.077; 0.053) | -0.034<br>(-0.100; 0.030) | 0.004<br>(-0.087; 0.095) |
| Tet VII | -0.009<br>(-0.098; 0.080) | **-0.074**<br>**(-0.146; -0.001)** | **-0.072**<br>**(-0.144; -0.001)** | -0.040<br>(-0.118; 0.039) | -0.018<br>(-0.096; 0.061) | **-0.082**<br>**(-0.157; -0.006)** | -0.050<br>(-0.136; 0.035) |
| Para | 0.053<br>(-0.041; 0.148) | 0.045<br>(-0.068; 0.158) | 0.059<br>(-0.043; 0.160) | -0.0009<br>(-0.092; 0.092) | 0.031<br>(-0.063; 0.128) | -0.038<br>(-0.130; 0.052) | -0.028<br>(-0.117; 0.062) |
| Blep | **0.126**<br>**(0.001; 0.249)** | -0.084<br>(-0.218; 0.050) | 0.040<br>(-0.126; 0.205) | 0.024<br>(-0.099; 0.144) | 0.043<br>(-0.085; 0.170) | -0.0003<br>(-0.157; 0.155) | -0.060<br>(-0.184; 0.067) |
| Eug | 0.016<br>(-0.129; 0.162) | -0.065<br>(-0.215; 0.084) | -0.013<br>(-0.170; 0.140) | -0.123<br>(-0.259; -0.008) | 0.083<br>(-0.047; 0.212) | 0.011<br>(-0137; 0.159) | 0.086<br>(-0.055; 0.225) |

Table S14: Statistical models for the multi-variate analysis of seven phenotypic traits. The analysis compares lines evolved at different selection temperatures (including evolutionary control) assayed at the other selection temperatures. Each row is a different model; the best model is highlighted in bold.

| Species | Model | Selection line random term | WAIC | Δ WAIC | SE | WAIC weight |
|---|---|---|---|---|---|---|
| Tet I | Intercept | Yes | 2796.14 | 9.5 | 143.56 | 0 |
| | **Selection temperature** | **Yes** | **2786.64** | | **140.33** | **1** |
| Tet VII | Intercept | Yes | 2672.60 | 28.55 | 83.51 | 0 |
| | **Selection temperature** | **Yes** | **2644.05** | | **84.59** | **1** |
| Para | Intercept | Yes | 4634.74 | 6.62 | 101.65 | 0 |
| | **Selection temperature** | **Yes** | **4628.12** | | **101.91** | **1** |
| Blep | Intercept | Yes | 1166.14 | 9.5 | 99.22 | 0.01 |
| | **Selection temperature** | **Yes** | **1156.64** | | **99.78** | **0.99** |
| Eug | Intercept | Yes | 1176.17 | 26.34 | 62.88 | 0 |
| | **Selection temperature** | **Yes** | **1149.83** | | **61.89** | **1** |

Table S15: Median slope and 95% compatibility intervals obtained from posterior distributions of the multi-variate analysis of 7 traits. The analysis compares lines evolved at different selection temperatures (including evolutionary control) assayed at the other selection temperatures. Slopes different from 0 are in bold.

| Species | $r_0$ | Max density | Bioarea | Major axis | Minor axis | Speed | Tortuosity |
|---|---|---|---|---|---|---|---|
| Tet I | -0.0005 (-0.024; 0.023) | -0.013 (-0.037; 0.009) | -0.029 (-0.052; 0.005) | 0.022 (-0.001; 0.045) | 0.015 (-0.007; 0.038) | -0.0008 (-0.022; 0.020) | 0.017 (-0.006; 0.041) |
| Tet VII | -0.018 (-0.042; 0.005) | 0.001 (-0.022; 0.025) | **-0.032 (-0.055; -0.009)** | 0.007 (-0.015; 0.031) | 0.013 (-0.009; 0.036) | **-0.055 (-0.076; -0.034)** | 0.005 (-0.017; 0.028) |
| Para | 0.001 (-0.017; 0.020) | **-0.028 (-0.046; -0.010)** | **-0.022 (-0.041; -0.004)** | **-0.025 (-0.043; -0.007)** | **-0.018 (-0.036; -0.0009)** | -0.003 (-0.021; 0.014) | **0.020 (0.001; 0.038)** |
| Blep | -0.025 (-0.069; 0.017) | -0.0006 (-0.047; 0.044) | **-0.054 (-0.096; -0.012)** | **-0.078 (-0.113; -0.042)** | **-0.057 (-0.098; -0.017)** | 0.033 (-0.007; 0.075) | **0.083 (0.046; 0.119)** |
| Eug | **-0.050 (-0.093; -0.007)** | **-0.048 (-0.092; -0.005)** | -0.022 (-0.067; 0.023) | **-0.066 (-0.109; -0.024)** | -0.033 (-0.071; 0.004) | **0.102 (0.068; 0.136)** | **0.105 (0.070; 0.140)** |
